# A novel engineered T7 RNA polymerase produces therapeutic mRNA without inflammatory dsRNA byproducts

**DOI:** 10.64898/2026.11.25.769297

**Authors:** Rebecca Leylek, Keiko Hokeness, Naryeong Kim, Brian Roper, Luladey Ayalew, Steffen Lippold, Carolina Galan, Julien Camperi, Emily Freund, Siri Tahtinen

## Abstract

The therapeutic application of in vitro transcribed (IVT) messenger RNA (mRNA) is constrained by double-stranded RNA (dsRNA) impurities generated by wild-type (WT) T7 RNA polymerase (RNAP). These impurities trigger innate immune responses via RIG-I and MDA5, suppressing protein translation. Conventional polymerase engineering often trades dsRNA reduction for lower transcription processivity. Here, we report engineered T7 RNAP variants that limit this trade-off. Physicochemical profiling identified variant G3, which suppresses high-molecular-weight aggregates and 3′-loopback hairpins while preserving full catalytic yield, high capping efficiency, and sequence processivity. In primary dendritic cells, N1-methylpseudouridine (N1psU)-modified mRNA produced by G3 yielded up to a three-fold increase in protein expression compared to WT T7, without inducing CD86 maturation. Systemic *in vivo* delivery in mice confirmed superior therapeutic expression and completely abrogated pro-inflammatory cytokine release. Functional knockouts confirmed that G3-transcribed mRNA evades cytosolic RIG-I/MDA5-mediated translational suppression by selectively eliminating inflammatory dsRNA byproducts. Together, G3 offers a superior, scalable manufacturing solution for advanced mRNA therapeutics.

## Introduction

The landscape of modern medicine is undergoing a paradigm shift driven by the rapid expansion of cell and gene therapies (CGTs). Beyond their pivotal role in infectious disease vaccines (1, 2), messenger RNA (mRNA) have emerged as a versatile and scalable platform to enable these next-generation therapeutics, including protein replacement therapies, cancer immunotherapies, ex vivo CAR-T cell engineering, and *in vivo* genome editing utilizing mRNA-encoded nucleases such as Cas9 (3, 4). The clinical success of these diverse applications relies on the capacity to manufacture pure, potent mRNA at scale. The current industrial standard for mRNA production is *in vitro* transcription (IVT), an enzymatic process catalyzed by bacteriophage T7 RNA polymerase (RNAP) (5–7). While T7 RNAP is favored for its high processivity and fidelity, it possesses inherent enzymatic liabilities that complicate downstream processing and compromise therapeutic efficacy (7).

During transcription, wild-type (WT) T7 RNAP generates product-related impurities, most notably abortive transcripts and double-stranded RNA (dsRNA) species (8, 9). T7 RNAP produces dsRNA predominantly through two mechanisms: (i) 3’-loopback extension, where the enzyme turns back on its own transcript to self-prime transcription, and (ii) promoter-independent antisense transcription, resulting from the enzyme re-binding full-length transcripts (7, 10). These dsRNA impurities act as potent agonists of cytosolic innate immune sensors, such as Retinoic Acid-Inducible Gene I (RIG-I) and Melanoma Differentiation-Associated Protein 5 (MDA5) (11). While incorporation of modified nucleosides like N1-methylpseudouridine (N1psU) or 5-methoxyuridine (5moU) successfully evades endosomal Toll-like receptor (TLR) activation, modified nucleosides alone can be insufficient to prevent cytosolic RIG-I/MDA5 sensing when residual dsRNA impurities are present (12). Active cytosolic sensing triggers pro-inflammatory cytokine cascades and induces Protein Kinase R (PKR)-mediated phosphorylation of eukaryotic initiation factor 2 alpha, leading to global translational arrest and silencing therapeutic RNA expression (13).

To mitigate these impurities, industrial workflows currently rely on downstream purification techniques, such as ion-pair reversed-phase liquid chromatography (IPRP-LC) or cellulose chromatography (9, 14). However, these polishing steps significantly reduce overall mRNA yield, increase manufacturing cost, and complicate process scale-up. Consequently, efforts have turned toward engineering RNA polymerases capable of suppressing dsRNA formation at the enzymatic source (15). Yet, active-site modifications frequently encounter a mechanistic trade-off known as the “binding affinity dilemma” (16). Mutations that lower template binding affinity effectively reduce dsRNA by preventing re-binding events, but they often diminish processivity, leading to an unacceptable surge in premature abortive transcripts and a reduction in yield (17). Uncoupling this trade-off requires precise active-site engineering that preserves full-length processivity while preventing non-templated extensions and antisense re-binding.

To address this challenge, we engineered a library of novel T7 RNAP variants (G1–G5), each bearing targeted single-point mutations designed to eliminate dsRNA generation while preserving catalytic throughput. To comprehensively evaluate these variants against the full spectrum of T7-derived product impurities, including abortive transcripts, high-molecular-weight non-covalent aggregates, covalent 3’-loopback hairpins, and 5’-capping heterogeneity, we established a rigorous, multi-attribute physicochemical profiling pipeline with functional assessments *in vitro* and *in vivo* (**Figure 1**). Through this screen, candidate variant G3 emerged as successfully uncoupling the binding affinity dilemma, maintaining high capping efficiency and processivity across most transcripts, while suppressing the formation of non-covalent aggregates and 3’-loopback species. Transcripts synthesized by G3 evaded the RIG-I/MDA5-mediated translational suppression. Additionally, systemic delivery of G3-transcribed N1psU-modified mRNA markedly enhanced protein expression while maintaining a naive cytokine baseline in mice, demonstrating that intrinsic enzymatic purity can streamline manufacturing and produce high-quality mRNA required for next-generation mRNA therapeutics.

**Figure 1.**
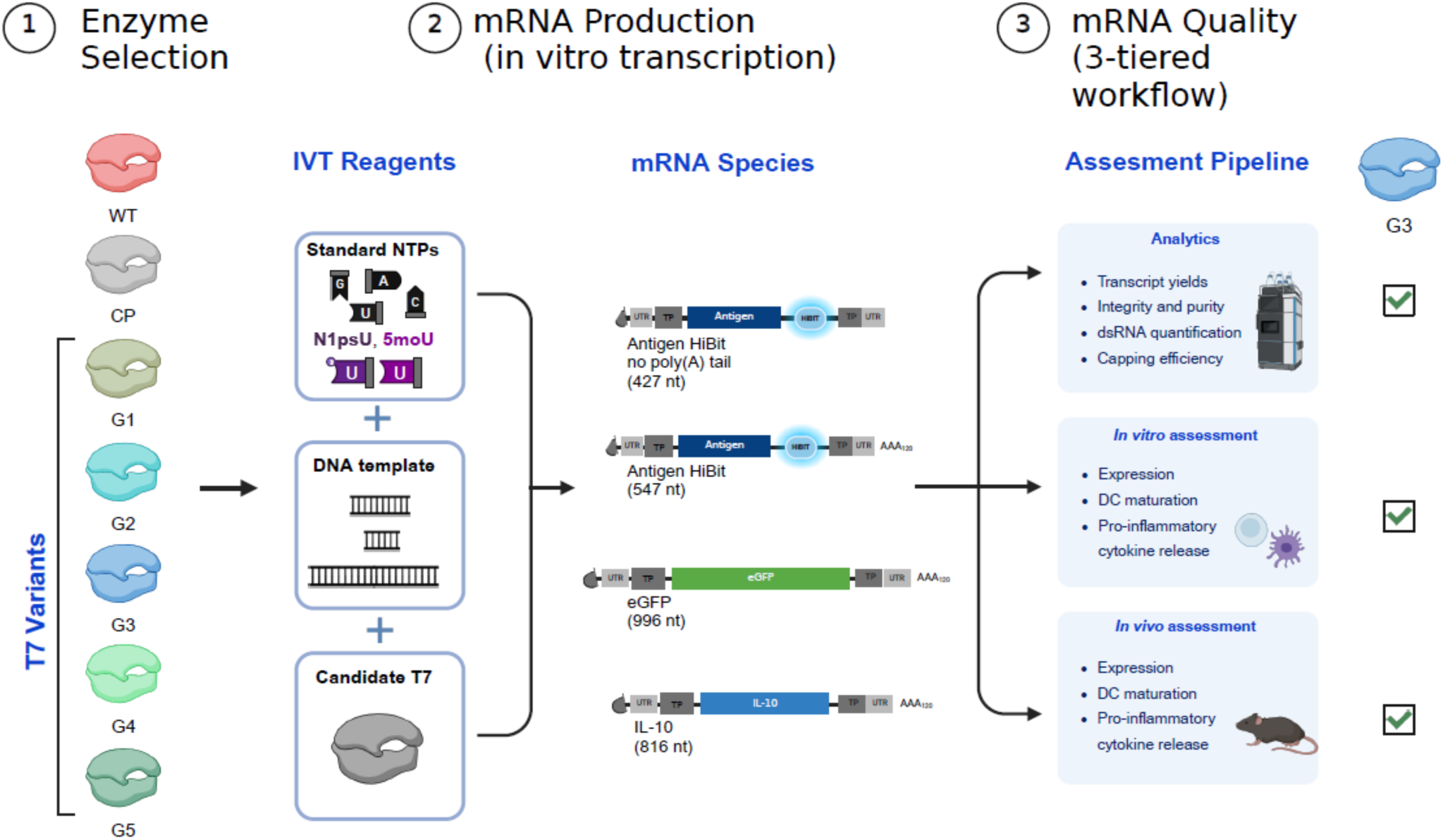
Experimental workflow for the synthesis and comprehensive evaluation of mRNA generated by engineered T7 RNA polymerase variants. Schematic depicting the 3-tiered pipeline used to characterize candidate T7 RNAP variants. (Left) In vitro transcription (IVT) is performed using a library of candidate T7 variants. The selected enzymes are combined with DNA templates of varying lengths and standard or modified nucleoside triphosphates (NTPs) to synthesize diverse mRNA constructs. (Right) The generated mRNA is subjected to a sequential 3-tiered evaluation workflow. The first tier consists of multi-attribute analytics to deeply profile physicochemical properties and impurity profiles. The second tier comprises *in vitro* functional assessments in cell models to evaluate protein expression efficiency (measured via percentage and gMFI), dendritic cell (DC) maturation, and pro-inflammatory cytokine release. The final tier utilizes *in vivo* murine models to validate systemic mRNA translation, DC maturation, and pro-inflammatory cytokine release.

## Results

### Physicochemical Profiling Identifies Engineered G3 and G4 Variants with Minimal dsRNA Formation and High Transcript Integrity

The enzymatic synthesis of mRNA via IVT involves a tightly coordinated catalytic cycle encompassing initiation, promoter clearance, processive elongation, and terminal release (18). Wild-type T7 RNAP is inherently prone to non-processivity, premature termination, and template switching, generating heterogeneous transcriptional byproducts including 5’-uncapped species, early abortive transcripts, poly(A) tail cleavage products, and structurally diverse dsRNA species. Because targeted active-site mutations alter enzyme processivity, promoter interaction, and terminal extension kinetics, engineered T7 RNAP variants inherently yield distinct impurity profiles (19). To systematically evaluate how variant engineering impacts key mRNA quality attributes without compromising overall synthesis yields, we designed three distinct mRNA constructs spanning diverse sequence architectures and lengths: a short antigen HiBIT-tagged construct lacking a poly(A) tail (427 nt), an antigen HiBIT-tagged construct containing a poly(A) tail (547 nt), and a reporter gene encoding enhanced green fluorescent protein (eGFP; 996 nt). We characterized transcripts from each class of RNA synthesized by engineered candidates G1–G5 and WT T7 RNAP and a commercial polymerase (CP) benchmark known for high fidelity and reduced dsRNA generation. Importantly, all engineered variants maintained overall IVT transcript yields comparable to WT T7 RNAP (∼3-5 mg/mL), confirming that impurity suppression was achieved without compromising catalytic throughput (**Supplementary Figure S1**).

First, we evaluated transcript length integrity and early termination using IP-RPLC (**Figure 2A**) and orthogonally confirmed via denaturing capillary gel electrophoresis (CGE) (**Supplementary Table S1**). IP-RPLC chromatograms obtained for eGFP constructs clearly resolved full-length main-peak mRNA from pre-peak impurities comprising short abortive transcripts, poly(A)-less species, and partial poly(A)-truncated intermediates (**Figure 2A**), as previously described (20). Quantification of relative peak area revealed that WT T7, CP, and variants G1–G4 maintained a high main-peak fraction with low pre-peak abortive levels (non-significant [*ns*] relative to WT T7) (**Figure 2A**). In contrast, variant G5 displayed a statistically significant accumulation of pre-peak species (****p < 0.0001), reflecting a propensity for premature termination due to a processivity trade-off (**Figure 2A**). Orthogonal capillary electrophoresis analysis confirmed these trends across all mRNA constructs (**Supplementary Table S1**), highlighting that candidates G3 and G4 maintained low abortive transcript levels, outperforming G2 and G5. Because the accumulation of pre-peak abortive species directly compromises translational capacity, these findings set an a priori expectation for reduced protein expression efficiency in downstream functional assays for these variants.

**Figure 2.**
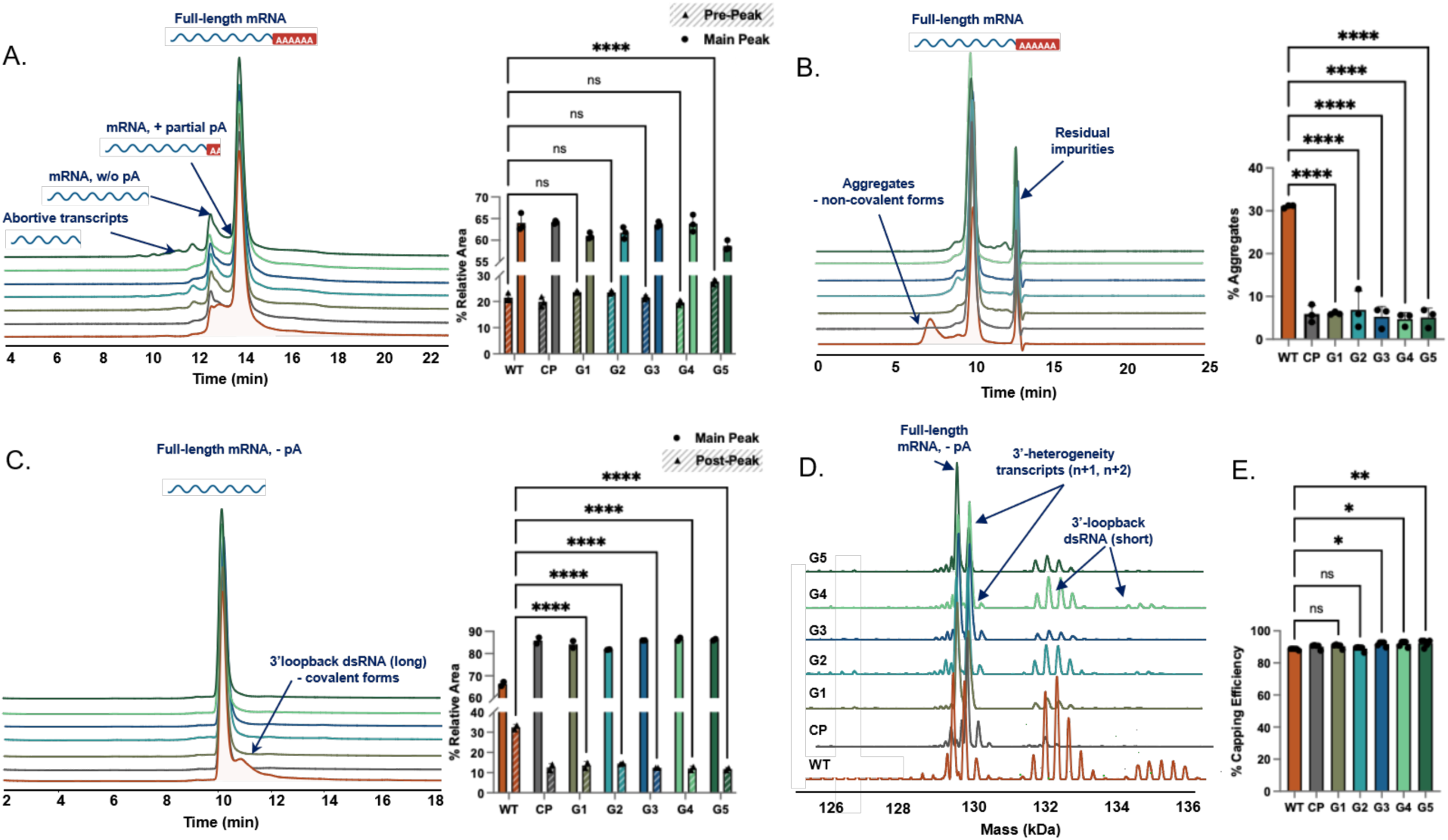
Multi-attribute physicochemical profiling and byproduct characterization of engineered T7 RNAP variants. **(A)** Denaturing IP-RPLC chromatograms (**left**) resolving full-length eGFP mRNA, partial poly(A), poly(A)-less species, and abortive transcripts, paired with a bar chart (**right**) quantifying the relative peak area percentage (% Pre-peak vs. % Main Peak) across T7 variants. **(B)** SEC chromatograms (**left**) resolving aggregates under non-denaturing conditions, paired with a bar chart (**right**) quantifying % aggregates across T7 variants for eGFP constructs. **(C)** IP-RPLC chromatograms of a poly(A)-less antigen HiBIT mRNA (**left**) resolving covalent long 3’-loopback dsRNA post-peaks, paired with a bar chart (**right**) quantifying the relative peak area percentage (% main peak vs. % post-peak). **(D)** Deconvoluted Native Mass Spectrometry spectra of poly(A)-less antigen HiBIT mRNA resolving 3’-heterogeneity (N+1, N+2) and short (<20 bp) covalent 3’-loopback dsRNA hairpins. **(E)** Bar chart quantifying % capping efficiency across variants, with individual sequence construct measurements overlaid as discrete points. Data are presented as mean +/- SD. For Panels A, B, and C, statistical analyses were performed across n = 3 independent in vitro transcription (IVT) reactions. For Panel E, statistical analyses were performed across n = 5 independent IVT reactions pooled across distinct sequence templates (antigen HiBiT without a poly(A) tail, antigen HiBiT with a poly(A) tail, and eGFP). Statistical significance was assessed via one-way ANOVA with Tukey’s post-hoc test (*ns* = not significant, *p < 0.05, **p < 0.01, ****p < 0.0001 relative to WT T7).

Having evaluated primary elongation processivity, we then deployed size-exclusion chromatography (SEC) and IP-RPLC to quantify higher-order secondary dsRNA byproducts (**Figure 2B–C**). Under non-denaturing conditions, SEC was used to quantify non-covalent aggregates, which include intermolecular sense-antisense dsRNA species (**Figure 2B**). WT T7 generated eGFP transcripts with a substantial aggregate burden (∼31.2%). In contrast, variants G1–G5 and CP dramatically suppressed non-covalent aggregate formation to ∼5–6% (****p < 0.0001 for all variants relative to WT T7) (**Figure 2B**). Similar suppression trends were obtained across all constructs, although the G2 variant exhibited slightly higher aggregate levels (**Supplementary Table S1**). To characterize covalent 3’-loopback dsRNA species, IP-RPLC analysis of the antigen HiBIT-tagged constructs lacking a poly(A) tail was performed to resolve post-peak species (**Figure 2C**). A poly(A)-less construct was intentionally selected to eliminate poly(A) tail length micro-heterogeneity, which causes chromatographic peak broadening. Therefore its removal maximizes chromatographic resolution and enables precise quantification of closely eluting 3’-loopback hairpins (20). While WT T7 exhibited a prominent post-peak tail corresponding to covalent 3’-loopback hairpins (∼31% post-peak area), all engineered variants (G1–G5) and CP maintained a flat post-peak baseline and a dominant main peak (>81–87% main peak area by IP-RPLC, ****p < 0.0001 relative to WT T7) (**Figure 2C**).

To orthogonally confirm the reduction of dsRNA species using a standard immunological assay, we performed anti-dsRNA antibody (J2) dot blot analysis on transcribed mRNAs. Consistent with our SEC and IP-RPLC profiling, commercial polymerase and engineered variants G1–G5 demonstrated a marked reduction in total dsRNA signal relative to WT T7 across evaluated constructs (**Supplementary Figure S2**). This qualitative immunochemical assessment directly mirrors the quantitative suppression of non-covalent aggregates and 3’-loopback hairpins observed in our chromatographic assays.

Conventional analytical techniques suffer from inherent physical and biochemical detection limits, specifically, IP-RPLC and CGE lack the size resolution required to resolve short (<20 bp) 3’-loopback hairpins from full-length linear transcripts, and the J2 anti-dsRNA antibody fails to recognize dsRNA helices shorter than 40 bp (21). To overcome this, we conducted native mass spectrometry (MS) analysis on the antigen HiBIT-tagged constructs lacking a poly(A) tail to resolve terminal 3’-end fine structure at single-nucleotide mass resolution (**Figure 2D**). The WT T7 spectrum displayed a characteristic multi-peak distribution trailing the main product, corresponding to non-templated nucleotide additions (N-1, N+2 3’-heterogeneity) and short (<20 bp) covalent 3’-loopback dsRNA hairpins caused by enzyme slippage (**Figure 2D**). Conversely, candidate variants, most prominently the CP benchmark, G1, and G3, produced highly homogeneous mass spectra dominated by a single major peak matching the theoretical molecular weight of the full-length transcript (**Figure 2D**).

Finally, to confirm that active-site mutations engineered to reduce dsRNA and abortive species did not compromise co-transcriptional capping efficiency, we assessed cap analog incorporation using site-specific RNase H cleavage coupled with IP-RPLC (**Figure 2E**). Across all evaluated mRNA architectures, all engineered candidate variants (G1–G5) generated mRNA species with high capping efficiencies ranging from ∼88% to ∼90%. Across distinct sequence templates, candidates G1–G5 exhibited capping performance comparable to or exceeding WT T7 (∼84.9–88.9%) and matching the CP benchmark (∼88.8–91.1%) (**Figure 2E**). While candidates G1, G2, and CP showed no statistically significant differences compared to WT T7, candidates G3, G4, and G5 achieved slightly higher, statistically significant capping increases (*p < 0.05p < 0.01) (**Figure 2E**). Individual sequence data points overlaid on the summary bars confirm low variance across templates, demonstrating that active-site modifications in G1–G5 do not adversely affect 5’-initiator recognition or cap analog incorporation regardless of sequence context (**Figure 2E**).

### G3 Mutants Produce Potent mRNA Without Inducing Innate Immune Activation *In Vitro*

To evaluate whether the improved purity and reduced dsRNA content of mRNA produced with the engineered T7 variants translates to enhanced functional performance, we assayed mRNA expression dynamics and innate immune activation *in vitro*. First, we synthesized unmodified (UTP) mRNA encoding either eGFP or a HiBIT-tagged antigen using WT T7, CP, and variants G1–G5. Each mRNA was delivered into C1R cells via electroporation, and expression was assessed by flow cytometry analysis for eGFP or by luminescence for HiBiT (**Figure 3A**). Transcripts of both target reporter species generated by all five candidate variants (G1-G5) produced significantly higher protein expression relative to WT T7. Specifically, G3 demonstrated the highest expression levels as assessed by eGFP geometric mean fluorescence intensity (gMFI) and HiBit luminescence readouts, matching or outperforming the commercial polymerase benchmark (**Figure 3B**).

**Figure 3:**
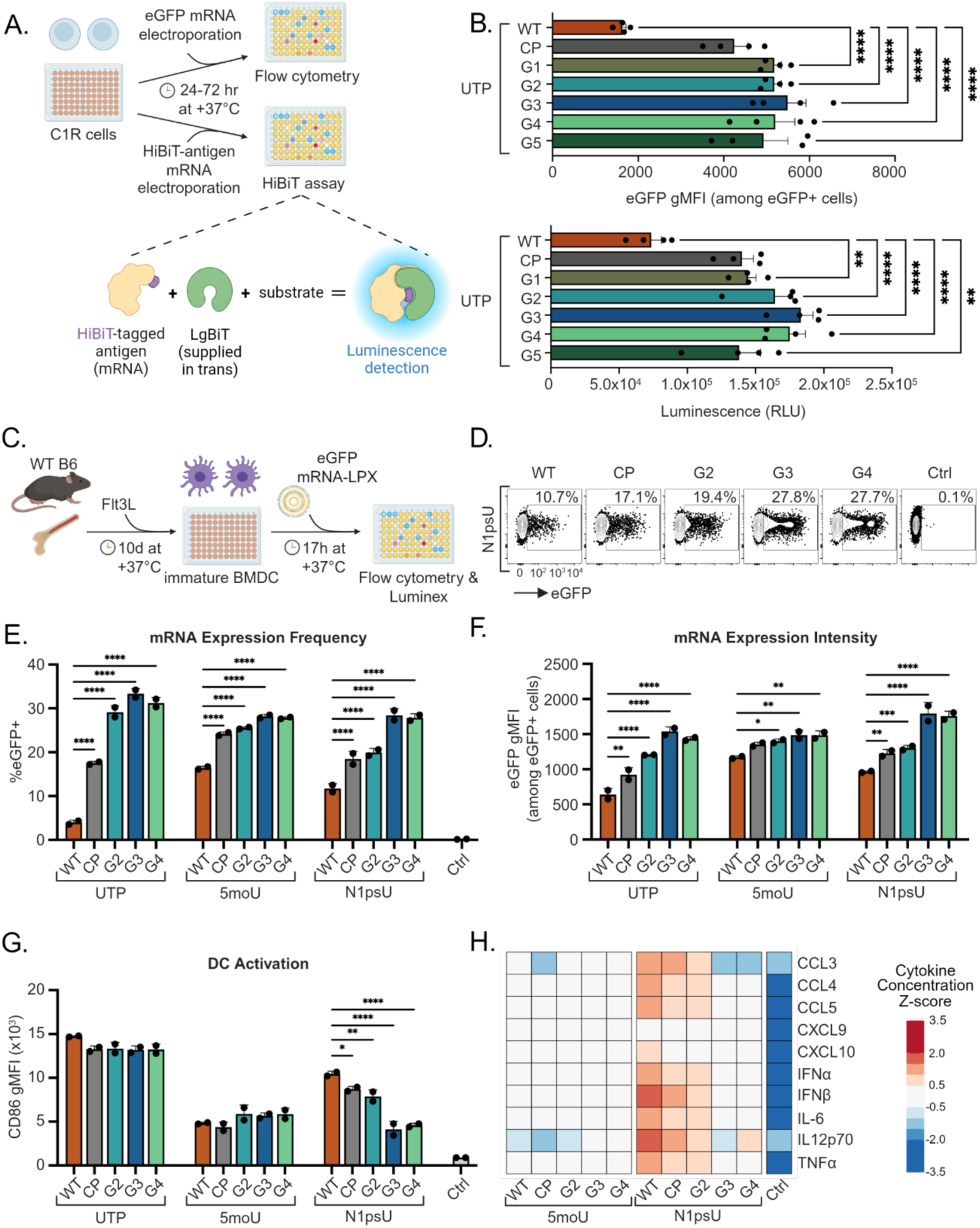
G3 mutants produce the potent mRNA without inducing innate immune stimulation *in vitro*. **(A)** Schematic depicting workflow to assess expression level of unmodified (UTP) mRNA encoding either eGFP or antigen-HiBiT electroporated into C1R cells. **(B)** Intensity of eGFP expression represented as geometric mean fluorescence intensity (gMFI) **(upper panel**) or luminescence-based measurement of HiBit expression (**lower panel**) in C1R cells (n=4 technical replicates). **(C)** Schematic depicting workflow to assess expression of LPX formulated UTP, 5moU or N1psU mRNA encoding eGFP in BMDCs. **(D)** Representative flow cytometry readouts of N1psU eGFP mRNA from BMDC assays (n=2 technical replicates). **(E)** Frequency of mRNA expressing cells (% eGFP+) among cDC2 BMDC population. **(F)** Intensity of expression represented as eGFP gMFI among eGFP+ cDC2 BMDCs. **(G)** DC activation represented by expression of the activation marker CD86 on BMDCs. (**H**) Cytokine secretion from BMDCs. Barplots show mean + standard error of mean (**B**) or mean + standard deviation (**E-G**). Statistics from one-way ANOVA with Tukey’s multiple comparisons test (**B**) or two-way ANOVA with Dunnett’s multiple comparisons test, tested against WT T7 for each uridine modification (**E-G**), *p<0.05, **p<0.01, ***p<0.001, ****p<0.0001.

To determine whether these translational gains extended to primary immune cells highly sensitive to dsRNA-mediated pattern recognition, we next evaluated functional performance in Flt3L-differentiated bone marrow-derived dendritic cells (BMDCs). Building upon our cell line screen, we investigated the compatibility of top-performing variants (G3 and G4, alongside candidate G2) across distinct nucleotide chemistries. Specifically, eGFP mRNAs, either UTP or modified with 5moU or N1psU, were synthesized, formulated into lipoplexes (mRNA-LPX), and incubated with BMDCs for 17 hours prior to flow cytometric analysis. (**Figure 3C**, **Supplementary Figure S3A**). Across all three uridine chemistries, mRNA produced by G2, G3, and G4 consistently resulted in higher percentages of eGFP-positive cells and elevated translation levels (eGFP gMFI) in conventional type 2 dendritic cells (cDC2s) compared to the WT T7 (**Figure 3D–F**). Notably, N1psU-modified mRNA generated by G3 yielded a nearly three-fold increase in eGFP+ cell frequency relative to WT T7 (**Figure 3D, E**). This robust enhancement in translation was similarly observed in the cDC1 population, where G3 and G4 variants drove superior mRNA expression frequency and intensity across all tested uridine modifications (**Supplementary Figure S3B, C**).

Because dsRNA contaminants act as potent agonists of pattern recognition receptors, we also assessed dendritic cell activation by measuring surface expression of the co-stimulatory marker CD86 (**Figure 3G**). As expected, BMDCs treated with mRNAs containing UTP showed high levels of surface CD86 regardless of the T7 enzyme used, likely due to TLR7 recognition of the unmodified uridine residues (22). In contrast, BMDCs treated with 5moU mRNAs showed similar, low levels of CD86 expression across all T7 enzyme conditions. Interestingly, BMDCs treated with N1psU-modified mRNAs exhibited profound differences depending on the T7 variants used for synthesis. WT T7- and CP-transcribed mRNAs induced pronounced dendritic cell activation. CD86 levels induced by delivery of G2-transcribed mRNA were significantly lower than those of WT T7, but remained higher than those of G3- and G4-transcribed mRNAs, which maintained a non-activating baseline closer to untreated controls (**Figure 3G**). Furthermore, G3 and G4-transcribed N1psU mRNA did not induce pro-inflammatory cytokine secretion in BMDCs, unlike mRNA generated by G2, WT T7 or the CP benchmark enzyme (**Figure 3H**). These findings suggest that the reduction of dsRNA byproducts in G3- and G4-derived transcripts prevents innate immune activation and enhances protein output in cells sensitive to dsRNA contaminants.

Integrating these *in vitro* results with our multi-attribute physical profiling, spanning length processivity (**Figure 2A** and **Supplementary Table S1**), aggregate suppression (**Figure 2B–C**), native MS 3’-homogeneity (**Figure 2D**), and capping efficiency (**Figure 2E**), we selected candidate G3 for further evaluation. Whereas G2 and G5 suffered from elevated abortive transcripts or residual innate immune activation, and G1 and G4 showed lower 3’-end homogeneity or reduced expression across reporter assays, G3 consistently delivered superior transcript homogeneity and dsRNA suppression without sacrificing yield or capping efficiency, achieving the highest *in vitro* potency across multiple coding sequences (CDSs).

### G3 maintains processivity across most 5’ UTRs but remains subject to sequence-dependent failures

Having established that the top candidate variant G3 exhibits minimal processivity issues across CDSs of varying lengths, we next evaluated its performance on untranslated regions (UTRs) of diverse structures. Because engineered UTRs often feature complex secondary structures and sequence motifs that challenge T7 RNA polymerase, we directly compared G3’s transcriptional performance against both WT and CP benchmarks. To evaluate how 5’-UTR sequence and structure impact transcriptional efficiency, we synthesized eGFP mRNA from 10 DNA templates containing distinct 5’ UTR sequences (UTRs 1–10) using WT T7, CP, and the top candidate variant G3 across three nucleotide conditions: unmodified UTP, 5moU, and N1psU (**Figure 4A-B**). We assessed the RNA yield to assay the throughput of each variant across each UTR. Across the majority of the tested templates, specifically UTRs 1, 3, 5, 6, 9, and 10, G3 generated robust mRNA yields (∼160 to 270 µg) comparable to both WT T7 and CP regardless of the uridine modification used (**Figure 4C**). However, specific UTRs revealed distinct variant-and nucleotide-dependent sensitivities. Notably, UTR-2 proved challenging exclusively for G3, which exhibited substantially reduced yields across all conditions (dropping to near-zero with N1psU), whereas both WT T7 and CP maintained high yields. Similarly, on UTR-7, both G3 and CP showed reduced yields relative to WT when incorporating modified nucleotides (5moU and N1psU), but under unmodified UTP conditions, only G3 failed to maintain high yields while CP performed comparably to WT. G3 also showed reduced efficiency on UTR-8 across all conditions, while UTR-4 proved universally challenging for all three polymerases, resulting in low yields across every condition and near-complete loss of transcription for G3 under UTP and N1psU **(Figure 4C).**

**Figure 4:**
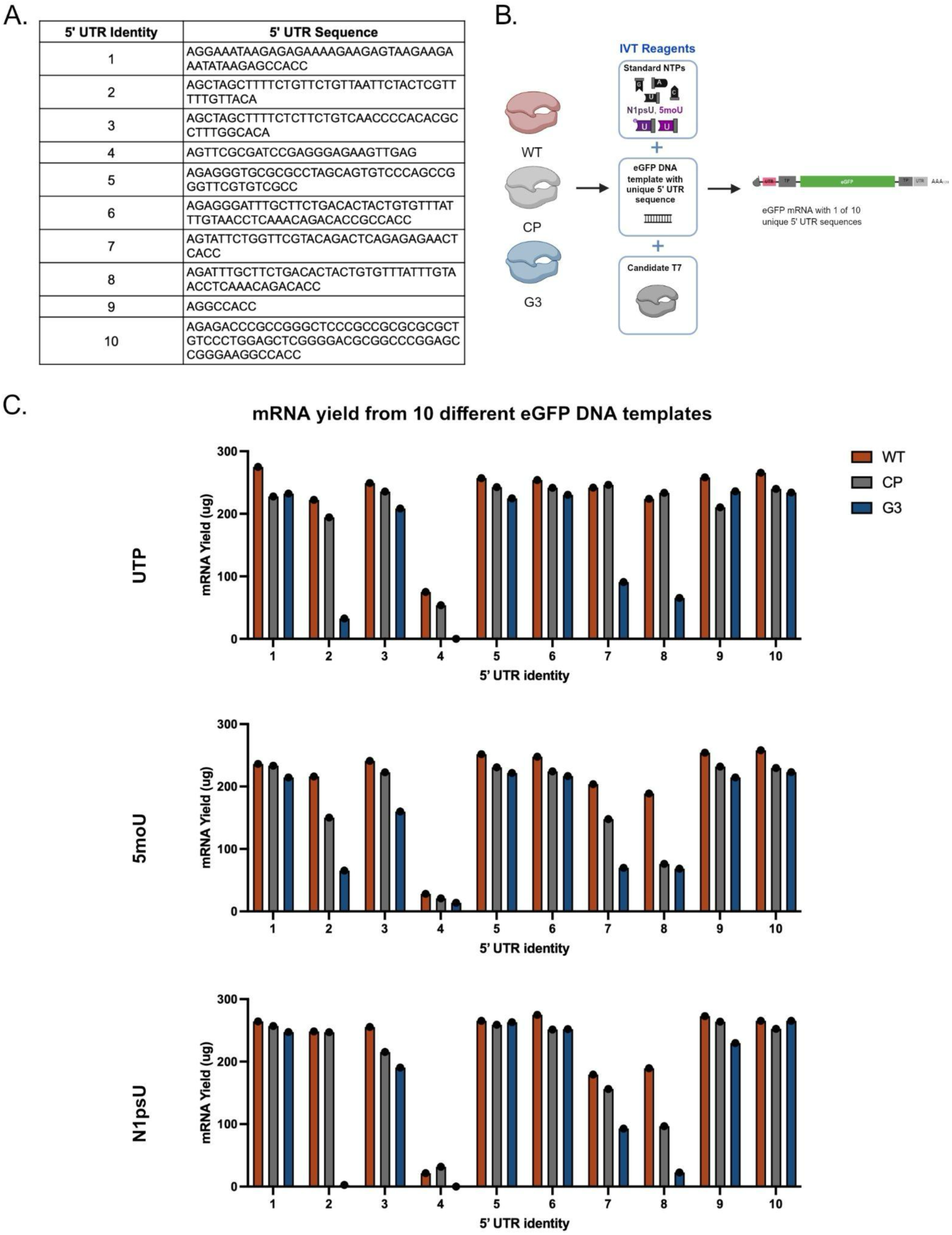
Characterizing processivity of WT T7, CP, and G3 across unique 5’ UTR sequences and nucleotide modifications. **(A)** Table containing the 10 distinct 5’ UTR sequences appended to the eGFP coding sequence. UTR-1 is the standard UTR sequence used in all other eGFP preps for subsequent *in vitro* and *in vivo* experiments. **(B)** Schematic of mRNA production with WT, CP, and G3 incorporating UTP, 5moU, or N1psU. Each IVT reaction used a DNA template with a different 5’ UTR sequence, paired with the eGFP coding sequence. **(C)** Comparison of total mRNA yield (in μg) for each IVT reaction produced with WT T7, CP, or G3 across eGFP DNA templates containing 1 of 10 unique 5’ UTR sequences. Bar graphs depict yields from IVT reactions incorporating unmodified UTP **(top)**, 5moU **(middle)**, and N1psU **(bottom)**.

Taken together, these findings demonstrate that the traditional binding affinity paradigm is not fully ameliorated; while G3 produces fewer IVT contaminants and concomitantly higher performance across diverse coding sequences, it remains more sensitive to challenging 5’ UTR sequences than its WT T7 and CP counterparts. Consequently, this highlights the critical need to pair the G3 candidate with compatible, optimal UTR sequences to achieve high transcriptional yields with superior mRNA purity, integrity, and overall performance. Nevertheless, G3 remains the variant capable of achieving the highest overall potency, prompting us to next verify whether this superior functional performance translates *in vivo*.

### High Protein Production and Minimal Innate Immune Activation with G3-Produced mRNA *In Vivo*

Having established that candidate G3 prevents dendritic cell activation *in vitro*, we next sought to determine whether this reduced innate immune activation translates to enhanced protein expression and reduced systemic inflammatory responses *in vivo*. Using an established systemic delivery approach (23, 24), mice were intravenously injected with 25 µg of eGFP-encoding mRNA formulated into LPX. Translation efficiency was assessed 24 hours post-injection across splenic antigen-presenting cell (APC) populations (**Figure 5A**, **Supplementary Figure S4A-B**). Systemic delivery of G3 transcribed N1psU-modified eGFP mRNA drove robust protein expression, achieving significantly higher eGFP gMFI in cDC2s and macrophages compared to WT T7-derived transcripts (**Figure 5B**). In contrast, no differences were observed in eGFP+ frequency in any of the analyzed splenic APC subtypes (**Supplementary Figure S4C**).

**Figure 5:**
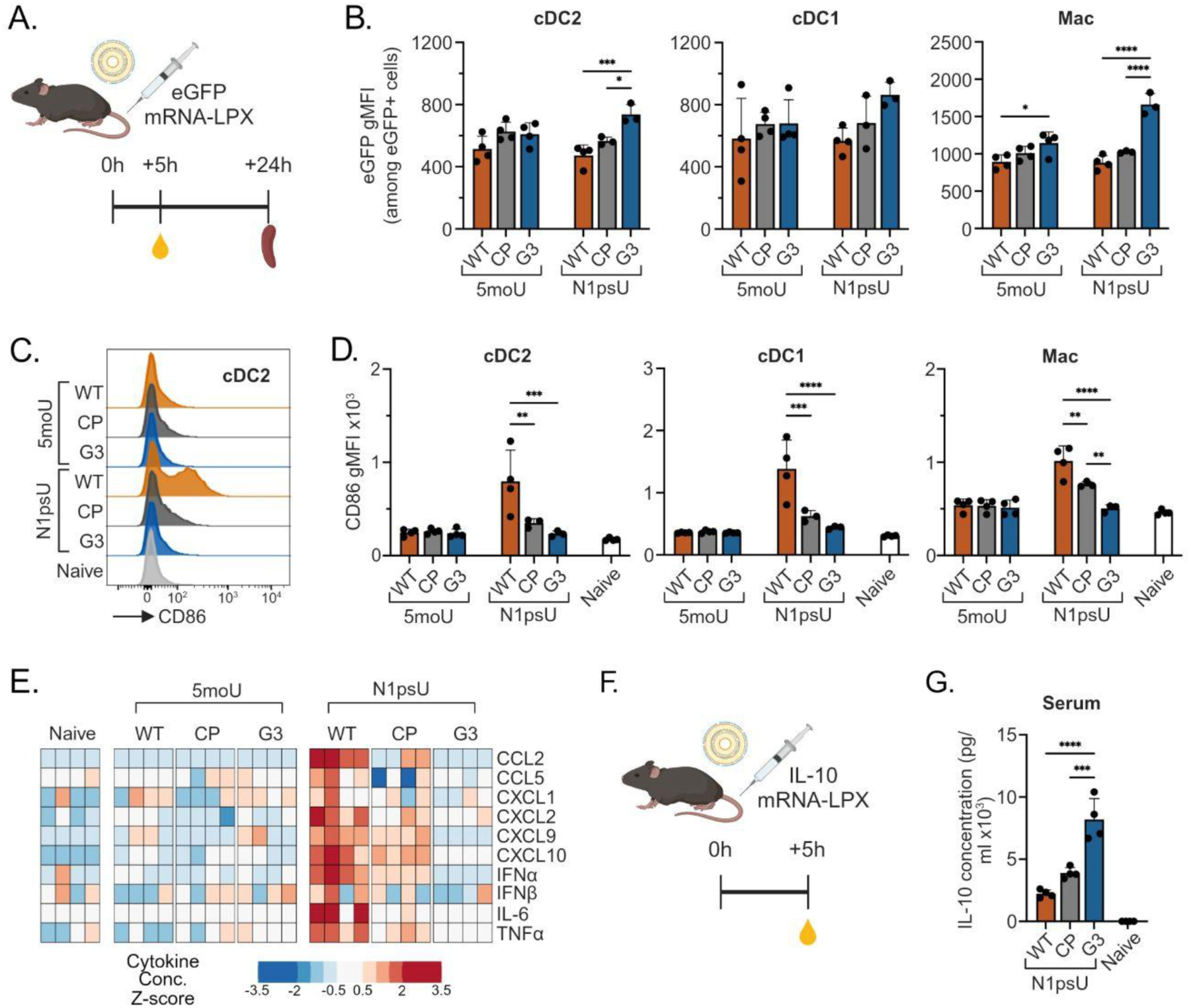
High protein production and minimal innate immune activation with G3 produced mRNA *in vivo*. **(A)** Schematic for *in vivo* eGFP mRNA experiment.25 µg eGFP mRNA-LPX was injected i.v. via the tail vein at 0h. Serum was collected at 5h for Luminex analysis. Spleens were collected at 24h for flow cytometry analysis (n=3-4 per group). **(B)** Intensity of expression represented as eGFP gMFI among eGFP+ splenic antigen-presenting cell populations. **(C)** Representative histograms for CD86 expression by cDC2s. **(D)** Antigen-presenting cell activation represented by CD86 expression on splenic populations. **(E)** Cytokines in serum measured by Luminex. **(F)** Schematic for *in vivo* IL-10 mRNA experiment.25 µg IL-10 mRNA-LPX was injected i.v. via the tail vein at 0h. Serum was collected at 5h for Luminex analysis. **(G)** IL-10 concentration in serum (n=4 per group). Barplots show mean + standard deviation. Statistics from two-way ANOVA with Tukey’s multiple comparisons test, *p<0.05, **p<0.01, ***p<0.001, ****p<0.0001.

We next sought to confirm our in vitro findings regarding the hierarchy of innate immune activation induced from the delivery of RNA produced by the WT, CP, and variant T7 enzymes. Flow cytometric analysis of the spleen revealed that WT T7-transcribed N1psU mRNA caused pronounced maturation of splenic cDC1, cDC2, and macrophage populations, as indicated by elevated surface expression of the co-stimulatory marker CD86 (**Figure 5C, D**). In stark contrast, CD86 expression in mice treated with G3-produced N1psU mRNA remained near baseline, closely mirroring naive controls. Consistent with our *in vitro* BMDC observations (**Figure 4H**), all mRNAs synthesized with 5moU induced negligible CD86 on splenic APCs, suggesting that different uridine analogs may influence how residual dsRNA impurities are formed during IVT and/or detected by cellular sensors in immune cells (25). Systemic inflammatory responses were further profiled by measuring serum cytokine concentrations at 5 hours post-injection (**Figure 5E**). WT T7-produced N1psU mRNA triggered a notable systemic release of pro-inflammatory cytokines, prominently including interferon-alpha (IFN-alpha), IFN-beta, interleukin 6 (IL-6), and tumor necrosis factor (TNF), alongside interferon-inducible chemokines C-X-C motif chemokine ligand 9 (CXCL9), CXCL10, and C-C motif chemokine ligand 2 (CCL2). Conversely, N1psU mRNA transcribed by the G3 variant completely abrogated this innate immune response, yielding cytokine profiles that were indistinguishable from naive baseline.

To confirm that circumventing innate immune sensing directly translates to enhanced production of a secreted therapeutic, we intravenously delivered RNA-LPX encoding the anti-inflammatory cytokine interleukin-10 (IL-10) (**Figure 5F**). In agreement with the fluorescent reporter data, mice receiving G3-transcribed N1psU IL-10 mRNA demonstrated superior therapeutic output. This group yielded significantly higher serum IL-10 concentrations compared to mice receiving mRNA synthesized by WT T7 or the commercial polymerase benchmark (**Figure 5G**). Collectively, these *in vivo* findings establish that G3 successfully eliminates inflammatory dsRNA contaminants at the source, thereby preventing translational suppression and maximizing therapeutic protein production.

### G3-Transcribed mRNA Evades RIG-I and MDA5-mediated Translational Suppression

To establish the precise molecular mechanism by which G3-transcribed mRNA evades cellular detection and translational suppression, we investigated the specific cytosolic dsRNA-sensing pathways responsible for innate immune responses to WT T7-transcribed mRNA. Product-related impurities such as dsRNA are potent agonists of cytosolic innate immune sensors, RIG-I and MDA5 (26). We utilized CRISPR-Cas9 ribonucleoproteins (RNPs) to engineer murine myeloid progenitor cells (27), generating distinct targeted knockouts (KO) for RIG-I or MDA5 alongside non-targeted controls (NTC). These edited progenitor cells were subsequently differentiated into immature BMDCs using Flt3L over 10 days (**Figure 6A**). Following differentiation, NTC, RIG-I KO, and MDA5 KO BMDCs were transfected with LPX-formulated eGFP mRNA synthesized by WT T7, the CP benchmark, or the G3 variant, and incubated for 17 hours. Translation efficiency was then evaluated by flow cytometry. For mRNA transcribed by WT T7, genetic ablation of either RIG-I or MDA5 resulted in a marked rescue of protein expression, calculated as the percentage increase over the NTC baseline (**Figure 6B-C**). Specifically, in the RIG-I KO and MDA5 KO cDC2 populations, WT T7 transcripts yielded approximately 40-50% increases in eGFP expression frequency, respectively, alongside robust 25–40% increases in overall expression intensity. This functional rescue indicates that residual dsRNA impurities present in WT transcripts actively trigger RIG-I and MDA5 to initiate translational suppression. In contrast, transfection with G3-transcribed mRNA yielded negligible increases in eGFP expression frequency (0–5%) in both knockout populations relative to the NTC baseline. While modest increases in expression intensity were observed (∼15%), they remained substantially lower than those triggered by WT T7. Because knocking out these primary dsRNA sensors provided minimal translational rescue for G3-derived transcripts, it confirms that G3-produced mRNA can inherently bypass the RIG-I and MDA5 signaling axes. By reducing inflammatory dsRNA byproducts during transcription, the engineered variant G3 prevents cytosolic sensor activation, proving that enzymatic purity at the transcript level directly drives functional innate immune evasion in primary myeloid cells.

**Figure 6:**
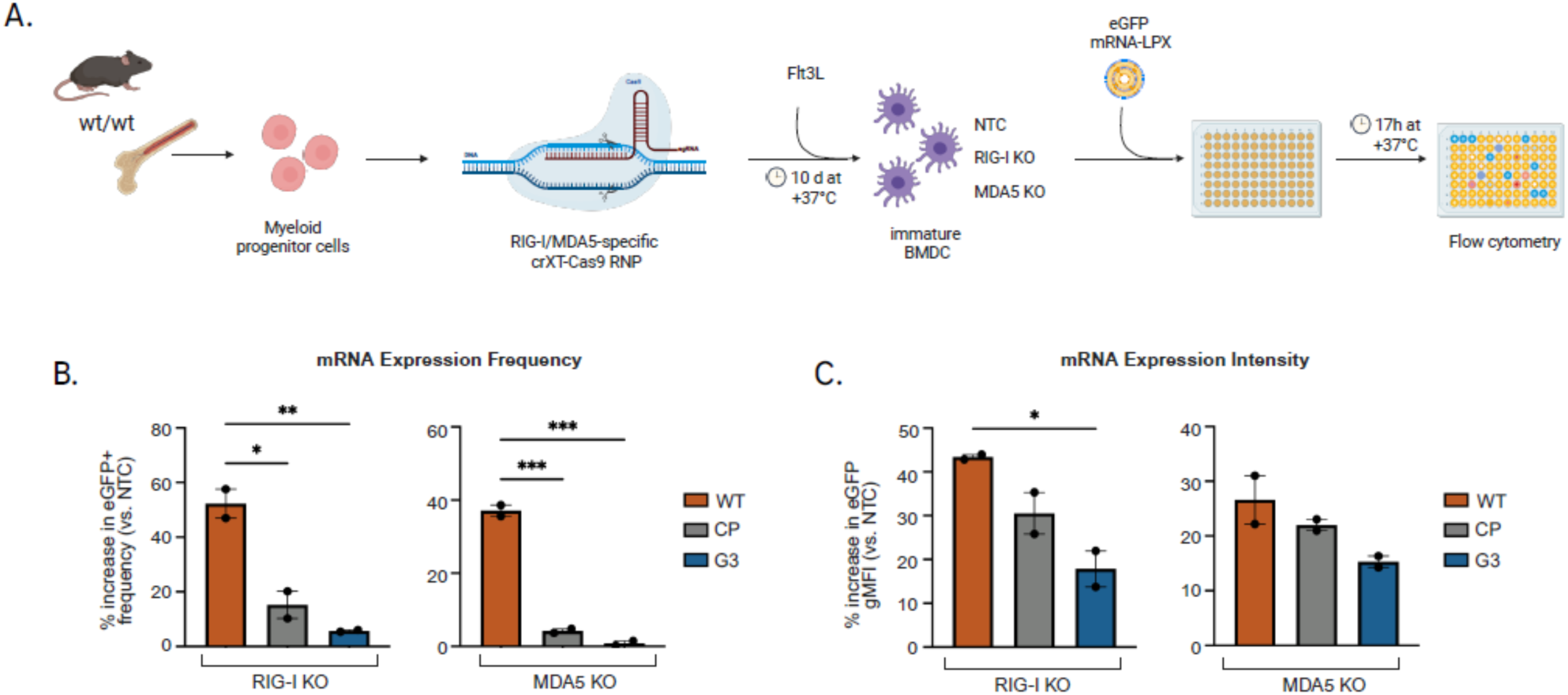
Genetic ablation of cytosolic sensors does not alter the translation of G3-produced mRNA. (**A**) Schematic workflow for generating RIG-I and MDA-5 knockout (KO) murine bone marrow-derived dendritic cells (BMDCs) and subsequent evaluation with N1psU mRNA-LPX. Murine myeloid progenitor cells were edited using CRISPR-Cas9 ribonucleoproteins (RNPs) targeting RIG-I or MDA-5, alongside non-targeted controls (NTC). (**B**) Percentage increase in mRNA expression frequency (% eGFP+) in the cDC2 population relative to the NTC baseline for each respective polymerase. (**C**) Percentage increase in translation intensity, represented as eGFP geometric mean fluorescence intensity (gMFI) among eGFP+ cDC2 BMDCs, relative to the NTC baseline for each respective polymerase. Barplots show mean + standard error of mean. Statistics from one-way ANOVA with Tukey’s multiple comparisons test, *p<0.05, **p<0.01.

## Discussion

In vitro transcription using bacteriophage T7 RNA polymerase is the cornerstone of mRNA manufacturing. However, the generation of product-related impurities, primarily dsRNA and abortive transcripts, remains a formidable bottleneck for the clinical translation of mRNA therapeutics, spanning mRNA vaccines as well as cell and gene therapies. A fundamental barrier in optimizing T7 RNAP is the “binding affinity dilemma.” Modifications that lower the enzyme’s binding affinity to the template often successfully prevent the self-priming and re-binding events that drive dsRNA formation, but they concurrently decrease processivity, leading to an unacceptable accumulation of premature abortive transcripts and reduced yield. In this study, we demonstrate that the engineered G3 variant successfully uncouples this trade-off for multiple CDS sequences and a majority of UTRs assayed. Our physicochemical profiling demonstrates that G3 maintains high transcript integrity and robust co-transcriptional capping efficiency (∼90%) while profoundly suppressing both non-covalent antisense aggregates and covalent 3’-extended loopback dsRNA structures across three mRNA constructs.

The performance of G3 represents a significant advancement over WT T7 and aligns with the recent engineering efforts reported in the literature (7, 19). Older strategies to mitigate dsRNA have largely relied on optimizing process conditions, such as adjusting magnesium concentrations or utilizing computationally designed thermotolerant T7 variants to run transcription at elevated temperatures (e.g., 48°C), which thermodynamically destabilize spurious RNA-template interactions (28, 29). Recent structural engineering approaches have focused on directly mutating the enzyme to reduce its RNA-dependent RNA polymerase and terminal transferase activities, such as the recently described combinatorial mutant M17 (30) or the T7-68 variant (31), alongside other newly reported engineered polymerases designed to enhance transcript purity (19). In addition, while previous computational engineering efforts have yielded T7 double mutants that successfully reduce immunostimulatory dsRNA to streamline manufacturing (7), such modifications can risk compromising overall enzyme processivity. Our data demonstrate that while experimental variants like G2 and G5 fell into the low-affinity trap, exhibiting abortive transcript impurities exceeding 23%, G3 achieved an abortive transcript profile comparable to the WT enzyme (∼20% for eGFP). This effect was even more pronounced for antigen HiBIT-tagged constructs, thereby ensuring full-length sequence fidelity alongside its significant dsRNA reduction.

The elimination of dsRNA at the enzymatic source is critical for maximizing the functional efficacy and required manufacturing ease for mRNA therapeutics. During cellular delivery, dsRNA acts as a potent pathogen-associated molecular pattern (PAMP). As our CRISPR-Cas9 knockout data confirm, WT T7-derived N1psU mRNA powerfully activates the RIG-I and MDA5 cytosolic sensors, signaling through the MAVS adaptor to induce a robust type I interferon (IFN-I) response and the release of pro-inflammatory chemokines such as CXCL9, CXCL10, and CCL2 (32). Beyond systemic reactogenicity, dsRNA contamination directly sabotages the primary function of the therapeutic mRNA: protein translation. Cytosolic dsRNA sensing and the induced IFN-I response activate the PKR pathway, which subsequently phosphorylates the alpha subunit of eukaryotic initiation factor 2 (eIF2α), leading to the rapid, global arrest of cellular translation (33). Our *in vitro* and *in vivo* studies demonstrate this dynamic: by minimizing dsRNA, G3-transcribed mRNA evades RIG-I/MDA5-mediated immune sensing and prevents translational shutdown, resulting in enhanced output of reporter proteins and therapeutic cytokines like IL-10.

The advantages of engineered T7 variants are particularly profound in the context of modern gene and cell therapies utilizing lipid formulations such as LPXs or lipid nanoparticles (LNPs). While transient immune activation and localized inflammation can serve as beneficial adjuvants for infectious disease or cancer vaccines (32, 34), they are severely detrimental to non-vaccine approaches. For applications requiring repeated systemic dosing (e.g., protein replacement therapies) or sustained high-level expression without triggering cellular apoptosis (e.g., CAR-T engineering or Cas9-mediated genome editing), bypassing this innate immune activation is essential to widen the therapeutic window. These advanced non-viral vectors are designed for highly efficient cytosolic delivery, which inadvertently ensures that any co-packaged dsRNA impurities are deposited directly adjacent to innate intracellular sensors. While the incorporation of modified nucleosides, such as N1psU or 5moU, effectively masks single-stranded mRNA from endosomal Toll-like receptors (22), our findings reiterate that nucleoside modification alone may not be sufficient to dampen inflammatory responses to mRNA generated by WT T7 RNAP. Only when synthesized with G3 did N1psU-modified transcripts maximize protein expression while maintaining a near-naive cytokine baseline *in vivo*. In contrast, 5moU incorporation showed prominent protein expression and low inflammatory responses even with WT T7, and similar incremental expression increases were not achieved with 5moU-modified mRNA using engineered T7 variants. This suggests that the specific biochemical nature of the uridine analog may dictate how residual dsRNA impurities influence the ultimate translational ceiling and inflammatory profile of therapeutic transcripts (25).

Ultimately, transitioning engineered T7 variants such as G3 into industrial IVT workflows presents a streamlined, highly scalable manufacturing paradigm. While current bioprocessing relies heavily on complex downstream purification techniques, such as cellulose chromatography or IP-RPLC, to partition dsRNA from the final drug product, our findings demonstrate that enzymatic suppression at the source can fundamentally reduce this dependency. By preventing antisense and loopback formation during synthesis, G3-mediated transcription simplifies downstream polishing pipelines, preserving overall mRNA yield and integrity while satisfying the stringent purity and safety profiles required for next-generation mRNA therapeutics.

## Materials and Methods

### mRNA Synthesis by In Vitro Transcription (IVT)

All mRNAs were synthesized from an initial DNA template produced by PCR as previously described (20). The templates were amplified from circularized plasmids with NEB’s Q5 Hot Start High-Fidelity 2X Master Mix and corresponding primers (600uL total). PCR products were purified with Invitrogen’s Dynabeads MyOne Carboxylic Acid beads. Nuclease-free water was used for the final elution. The purity of the purified DNA templates was assessed by e-Gel and Sanger Sequencing.

The following reagents were combined in a tube for the in vitro transcription reaction: 100mM CleanCapAG-3’OMe (4mM), 100mM ATP/CTP/GTP (5mM), 100mM UTP or N1psU or 5moU (5mM), 100mM DTT (4mM), 1M Mg(OAc)2 (10.5mM), 10X NEB Transcription Buffer (1X), 0.792U/uL Pyrophosphatase (0.002U/uL), and the DNA template at 0.025ug/uL. The WT T7 reaction received 40U/uL RiboLock RNase Inhibitor at 1U/uL and the 200U/uL WT T7 at 10U/uL, while the CP reaction received 20U/uL SUPERase-In at 1U/uL and 0.2mg/mL Cleanscribe RNA Polymerase (Trilink Biotechnologies, cat no. E-0107) at 0.016mg/mL. Each of the five Ginkgo mutant polymerase reactions received SUPERase-In at 1U/uL and 0.5mg/mL Ginkgo mutant T7 at 0.02mg/mL, as recommended by Ginkgo Bioworks. Each tube was brought up to a total of 200uL with Nuclease-Free Water and vortexed thoroughly. Tubes were quickly spun down, then incubated on a shaking ThermoMixer F2.0 (37C, 350rpm) for 2.5 hours. After the incubation, 5uL of 2U/uL TURBO DNase was added to each tube. Tubes were again vortexed and spun down before being returned to the ThermoMixer for an additional 15 minutes. Following the final incubation, 20uL of 500mM EDTA and 112.5uL of 7.5M LiCl were added to each tube. Tubes were frozen overnight, then spun down in a pre-cooled mini centrifuge (4C, 15min), producing a visible white pellet. The RNA pellet was washed with ice cold 70% ethanol twice and left to air dry for 10 minutes at RT. The pellet was resuspended with nuclease-free water until the RNA went into solution.

### IP-RPLC-UV Analysis

Chromatographic separations were conducted on an Acquity UPLC H-Class platform (Waters, Milford, MA) integrated with a quaternary pump, autosampler, and UV absorbance detector. Instrument control and data processing were carried out using Chromeleon software (version C.01.07 SR4 [505], Thermo Fisher Scientific, Santa Clara, CA). Messenger RNA species were resolved on a DNAPac RP column (2.1 mm I.D. × 100 mm, 4.0 μm particle size; Thermo Fisher Scientific). Eluent A consisted of an aqueous buffer containing 50 mM triethylamine (TEA) and 50 mM hexafluoroisopropanol (HFIP) adjusted to pH 9.3. Eluent B was an organic mixture of 25 mM TEA and 25 mM HFIP in 90% (v/v) methanol. The gradient profile was programmed as follows: initial hold at 5% B for 1 min, a linear increase from 5% to 18% B over 19 min, a rapid ramp to 70% B over 1 min, and a return to 5% B held for 5 min to re-equilibrate. The flow rate was set to 0.4 mL/min, the column compartment was heated to 80 °C, and optical detection was performed at a wavelength of 260 nm.

### SEC–UV Analysis

The size distribution of mRNA samples was evaluated using an SRT(R) SEC-1000 column (4.6 mm I.D. × 300 mm, 5 μm, 1000 Å pore size, stainless steel; Sepax Technologies, Inc., Newark, DE). Isocratic elution was carried out using a mobile phase composed of 100 mM Tris–HCl and 300 mM NaCl (pH 7.5) at a flow rate of 0.350 mL/min. The column temperature was controlled at 25 °C, and injections of 10 μL were monitored via a diode array detector monitoring at 260 nm.

### Intact mRNA Analysis by Native Mass Spectrometry (nMS)

Intact mRNA characterization was carried out on an Orbitrap Q Exactive UHMR mass spectrometer featuring a nanospray flex ion source (Thermo Fisher Scientific, Santa Clara, CA). Samples were buffer-exchanged into 200 mM ammonium acetate using 10– 50 kDa molecular weight cut-off centrifugal filters (Millipore Sigma, MA). For poly(A)-tailed mRNA targets presenting highly complex, heterogeneous charge state distributions, 100 mM triethylammonium acetate was supplemented post-desalting to promote charge reduction and simplify the resulting mass spectra. Sample preparation involved diluting the analyte with 200 mM ammonium acetate to a total volume of 500 μL, performing five sequential wash cycles (centrifuged at 14, 000 × g for 5 min each), and recovering the concentrated retentate in a final volume of 20–50 μL (targeting a concentration range of 0.1–1 μg/μL). A 15 μL aliquot of the desalted sample was loaded into a borosilicate glass emitter for nanoESI infusion.

### 5’ Capping by RNase H digestion and IP-RPLC analysis

Evaluation of the 5’ cap structure commenced by annealing the sample to a complementary biotinylated DNA/RNA hybrid probe targeting the initial 18 nucleotides at the 5’ terminus of the mRNA construct. The target hybrid region was selectively cleaved using Ribonuclease H (RNase H), followed by isolation of the resulting fragments using streptavidin-coated magnetic beads (Invitrogen, cat. no.65001). Separation of the short capped and uncapped oligonucleotide cleavage products was carried out on an Acquity UPLC H-Class system (Waters) utilizing a DNAPac Reversed Phase column (2.1 mm I.D. × 100 mm, 4 µm; Thermo Fisher Scientific, Waltham, MA). Eluent A was prepared as 10 mM triethylamine and 50 mM HFIP in water at pH 9.0, while Eluent B was composed of 5 mM triethylamine and 25 mM HFIP in 90% methanol. Chromatographic separation was achieved using a gradient that held at 5% B for 1 min, increased linearly to 26% B over 20 min, ramped to 70% B over 2 min to wash the column, and finally returned to 5% B over 5 min for re-equilibration. The total method run time was 28 min with an injection volume of 5 μL, a column temperature maintained at 80 °C, and UV detection recorded at 260 nm using a diode array detector.

### Cell lines and electroporation

Assessment of mRNA expression was performed in an engineered human B cell line, C1R-T2A (35).5e6 cells per condition were harvested and washed one with PBS before gentle resuspension in 20uL of the Lonza SE Cell Line Nucleofector Solution. The HiBiT-encoding antigen and eGFP mRNAs were each made with the seven different polymerases and tested in duplicates.3ug of each mRNA was aliquoted and 5e6 cells were added to each mRNA aliquot. Untreated cells were also included as a control. The mixture was slowly transferred to a 16-well Nucleocuvette to avoid air bubbles, and electroporated with the program code CM-138. Cuvettes were immediately transferred to the incubator for 15 minutes before 100uL of pre-warmed media (IMDM + 20% FBS) was added to each well. The cuvettes were incubated for another 15 minutes to allow for cell recovery before the cells were transferred to 48-well plates containing 2mL of pre-warmed media. The 2mL of cells were then split between two plates for use at two different timepoints. Cells were cultured at +37°C with 5% CO_2_ for 24-48 hours. After incubation, cells treated with HiBiT-encoded antigen mRNA were prepared for a HiBiT luminescence assay while cells treated with eGFP were stained for flow cytometry.

### HiBiT luminescence measurement

Expression of HiBiT protein was analyzed with Promega’s Nano-Glo HiBiT Extracellular Detection System kit following manufacturer’s protocol. The 1mL of treated cells were transferred to a 96-well V-bottom plate in triplicates with 500k cells per well (200uL per well). The plate was centrifuged (300 x g, 5 minutes) and the supernatant was removed. The HiBiT reagent was prepared by adding the LgBiT protein (1:100) and Nano-Glo HiBiT Extracellular Substrate (1:50) to the Nano-Glo HiBIT Extracellular Buffer. Cells were resuspended in 50uL PBS, followed by the addition of an equal volume of the HiBiT solution to each well. The solution was mixed thoroughly and the plate was left to incubate in the dark at RT for 10 minutes. Following the incubation, the cell solution was slowly transferred to a clear-bottom, white opaque-wall 96-well plate to eliminate luminescence leaking into neighboring wells. The luminescence was measured with BioTek’s Synergy H1 microplate reader. To quantify HiBiT luminescence level, unused well measurements were averaged and subtracted from sample wells to account for background luminescence.

### Preparation of mRNA-LPX

RNA was formulated with liposomes consisting of DOTMA and DOPE at a charge ratio (+):(−) of 1.3:2, yielding negatively charged mRNA-LPX (24).

### Mice

All mice were used in accordance with protocols approved by the Genentech Institutional Animal Care and Use Committee and respected the principles of animal experimentation. C57BL/6J female mice at 6-10 weeks old were purchased from Jackson West. All mice had *ad libitum* access to standard chow and water. All animals were maintained under a 12-hrs light/dark cycle at 22 °C. All animal studies were reviewed and approved by Genentech’s Institutional Animal Care and Use Committee.

### Preparation of bone marrow-derived dendritic cells

Bone marrow-derived dendritic cells (BMDCs) were following the protocol with Flt3L published by Lutz et al (36). Briefly, femurs and tibias were dissected from two mice and sterilized by placing them in 70% ethanol for 4 minutes, then rinsed with phosphate-buffered saline (PBS). Bone ends were cut off, and bone marrow cells were flushed using a 25G needle and 10 mL syringe filled with RPMI 1640 without serum. A single-cell suspension was created by mechanically dissociating cell clumps and passing the solution through a 70 μm filter. The cells were centrifuged (400 x g for 5 minutes at room temperature), then resuspended in 3 mL ACK lysis buffer and allowed to incubate for 3 minutes at room temperature. Lysis was stopped by adding 10 mL of R10 media (RPMI 1640 with 10% heat inactivated FBS, 1X pen/strep, 1X glutamax or L-glutamine, and 55 mM beta-mercaptoethanol), and the cells were centrifuged again. Cells were counted using a ViCell-Blu and resuspended at a final concentration of 2e6 cells / mL in R10 media with 100 µg/mL murine Flt3L (Peprotech cat# 250-31L). Cells were plated with 5 mL per well in tissue-culture treated 6-well plates (Corning cat# 3516) and cultured in an incubator (37°C, 5% CO2) for 10 days with cytokine-containing media changed 1-2x. On day 10, non-adherent BMDCs were gently collected for use in assays.

### RNA Expression and cytokine-release in BMDCs

BMDC assay was performed as previously described (9). Briefly, BMDCs were counted and resuspended in fresh R10 media with 100 μg/mL of murine FLT3L and plated at a concentration of 200k cells in 180 μL per well of a 96-well round-bottom plate (Corning cat# 3799). mRNA-LPX were added to cells in the 96-well plate at a final concentration of 5 µg/ml mRNA and pipetted up and down to mix. Cells were placed in an incubator (+37°C, 5% CO_2_) overnight, ∼16h. The next day, plates were centrifuged (700g for 4 min at +4°C).

### *In vivo* experiments and tissue processing

For *in vivo* experiments, 25 μg mRNA-LPX diluted in 150 mM saline was injected intravenously into C57BL/6J mice. Five hours after injection, mice were anesthetized with isoflurane and blood was collected retro-orbitally for serum cytokine analysis. For flow cytometry, spleens were collected in cold PBS, and single-cell suspensions were generated by mashing the spleen tissue with a syringe plunger, incubating for 15 minutes at room temperature in Hank’s-based Cell Dissociation Buffer (Gibco) with 12.5 µg/ml Liberase TL (Roche) and 50 µg/ml DNase I (ThermoFisher), then passing through a 70-μm cell strainer (BD Falcon). Splenocytes were washed with MACS buffer (1X PBS + 0.5% BSA + 2 mM EDTA) and centrifuged (500g for 5 minutes at room temperature), then resuspended in ACK lysis buffer (Gibco) for 8 minutes at room temperature to lyse red blood cells. MACS buffer was added to quench, and splenocytes were centrifuged again before proceeding to staining for flow cytometry.

### Luminex analysis

Cell culture supernatants or sera were collected and stored at −70 °C until Luminex analysis. Luminex was performed with a custom MILLIPLEX mouse panel 1 multiplex (Millipore Sigma), collected on a Flex-Map 3D (Luminex Corporation), and analyzed with Bio-Plex Manager.

### Flow cytometry

Cells were washed twice with PBS, then stained with LIVE/DEAD Fixable Blue (Invitrogen cat# L23105) or LIVE/DEAD Fixable Near-IR Dead Cell Stain kit (Invitrogen cat# L34975). In experiments with antibody stainings, cells were first incubated with TruStain FcX block (Biolegend cat# 101320) for 10 minutes at room temperature. For FACS analysis, cells were then washed with PBS and stained with an antibody cocktail (see **Supplementary Table S2**) for 20 minutes on ice. Cells were analyzed on a BD FACSymphony flow cytometer (BD Biosciences). Data analysis was performed with FlowJo software. Heat-killed cells and unstained cells were used as controls for live-dead and positive antibody staining.

### BMDC CRISPR Knockouts

Bone marrow was isolated as described above. On day 0, knockouts were performed as previously described (27). Briefly, TruCut V2 Cas9 (5 mg/ml; Thermo Fisher Scientific) was combined with 100 µM sgRNA at a 1:1 volume ratio and incubated for at least 10 minutes at room temperature to allow ribonucleoprotein (RNP) complexes to form. Two guides were used for each KO condition (see **Supplementary Table S3** for sequences). Bone marrow (BM) cells were washed twice with PBS. For each nucleofection reaction, 4e6 cells in 20 µl of P3 buffer from the P3 Primary Cell 4D-Nucleofector X Kit S (Lonza Bioscience) were combined with 8 µl of RNPs and pipet into one well of a nucleocuvette strip. Cells were nucleofected with a 4D Nucleofector X (Lonza Biosciences) with program CM-137. Warm media was added immediately; cells were then plated in 6-well plates and differentiated as above.

### Statistical analyses and data presentation

Statistical analyses and graphing were performed using GraphPad Prism v.11 for Mac OS, R software, and RStudio. Illustrations were created with BioRender. com. All results are expressed as mean ± s.e.m. or mean ± s.d. as indicated. One-way or two-way analysis of variance (ANOVA) was performed when more than two groups were compared, and multiple comparisons were corrected using Dunnett’s post hoc test or Tukey’s post hoc test. *P ≤ 0.05, **P ≤ 0.01, ***P ≤ 0.001 and ****P ≤ 0.0001. No statistical methods were used to predetermine sample size for animal experiments.

