## Supplemental Figures and Tables for "A novel engineered T7 RNA polymerase produces therapeutic mRNA without inflammatory dsRNA byproducts"

### Supplementary Figures

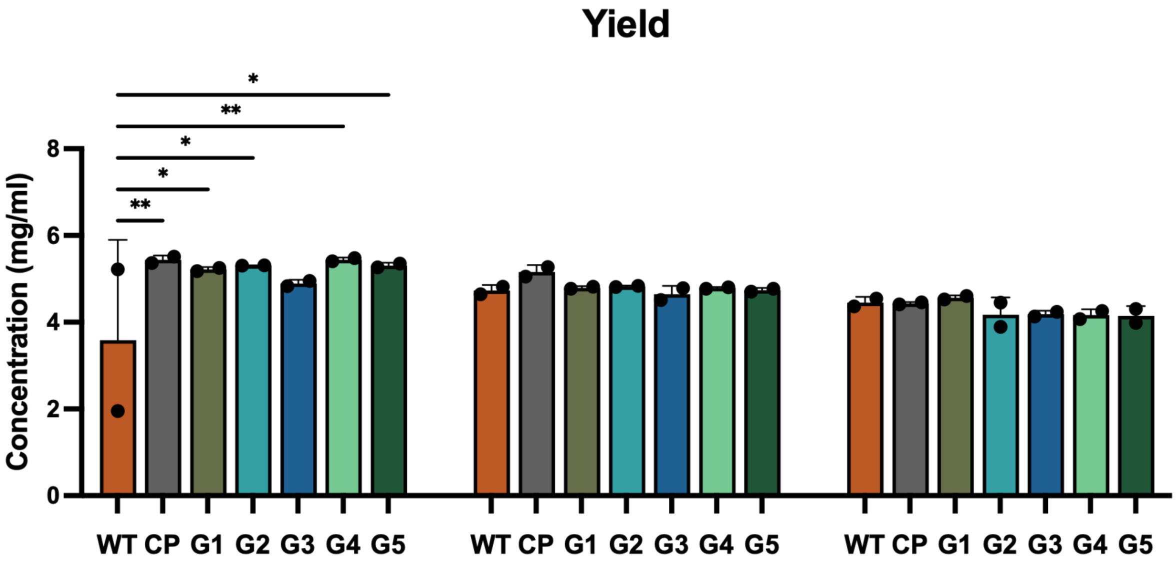


**Supplementary Figure S1.** Comparison of RNA yield (RNA concentration in mg/mL) from IVT reactions producing eGFP mRNA using WT, CP and all 5 mutant Gingko T7 RNA polymerases. Individual replicate data points (n=2) are displayed, with horizontal lines indicating the mean concentration for each enzyme treatment. Statistics from two-way ANOVA with Dunnett's multiple comparisons test, tested against WT T7 for each uridine modification, *p<0.05, **p<0.01.

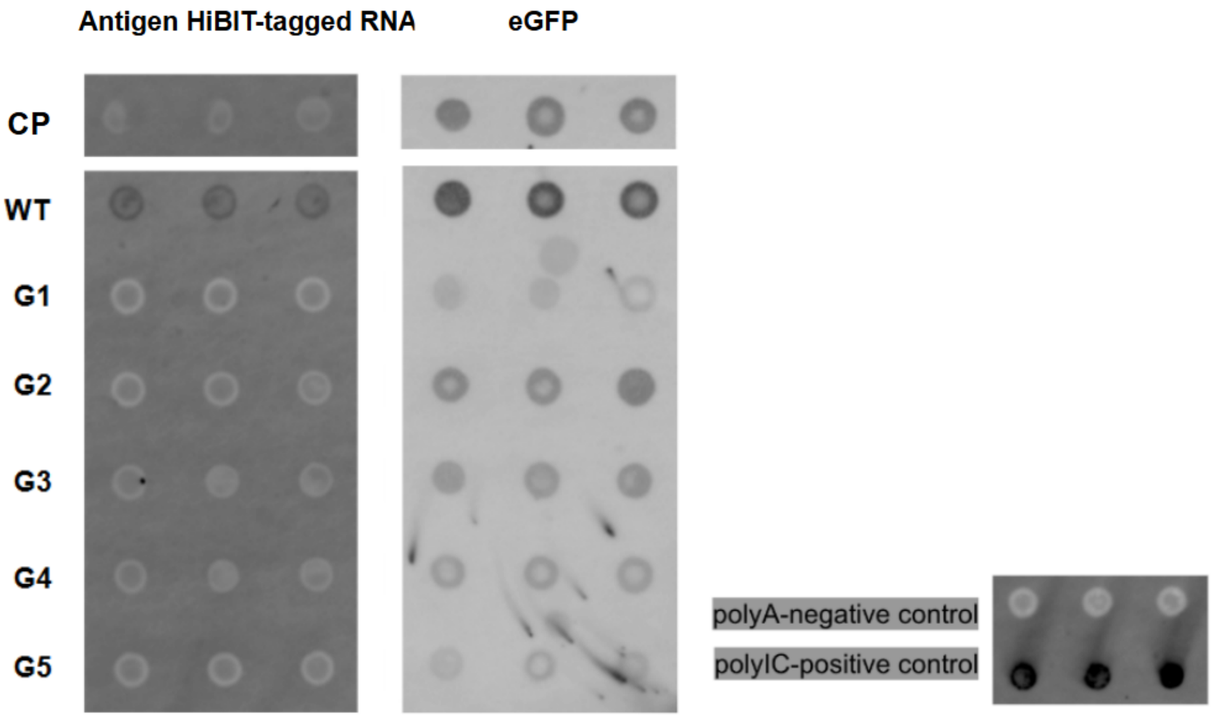


**Supplementary Figure S2.** Markedly Reduced dsRNA Contamination Across Mutant T7-Transcribed mRNA Constructs. Representative anti-dsRNA J2 dot blot images for two distinct mRNA constructs synthesized by wild-type (WT) T7 RNAP, commercial polymerase (CP), and engineered variants G1–G5. Transcripts evaluated include a poly(A)-less antigen HiBIT-tagged construct (427 nt) and an eGFP reporter mRNA (996 nt). Strong, persistent J2 immunoreactivity was observed for all WT T7-transcribed constructs, whereas signal intensity was markedly suppressed across all G-series variants and CP, regardless of transcript length or poly(A) tail status.

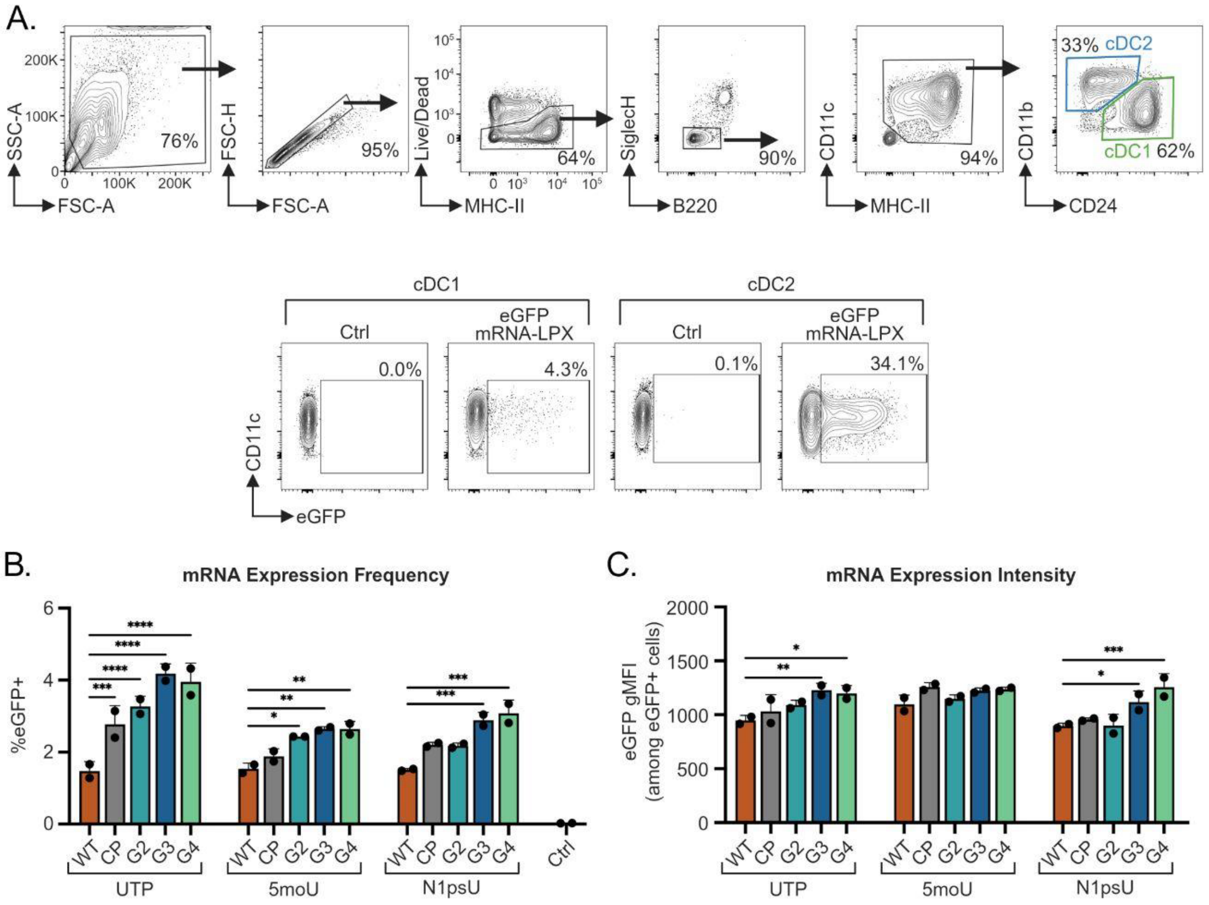


**Supplementary Figure S3. (A)** Representative flow cytometry gating strategy for BMDCs (top) and representative gates for eGFP+ cDC1 and cDC2 (bottom). **(B)** Frequency of mRNA-expressing cells (% eGFP+) among cDC1 BMDC population. **(C)** Intensity of expression represented as eGFP gMFI among eGFP+ cDC1 BMDCs. Barplots show mean + standard deviation. Statistics from two-way ANOVA with Dunnett's multiple comparisons test, tested against WT T7 for each uridine modification, *p<0.05, **p<0.01, ***p<0.001, ****p<0.0001.

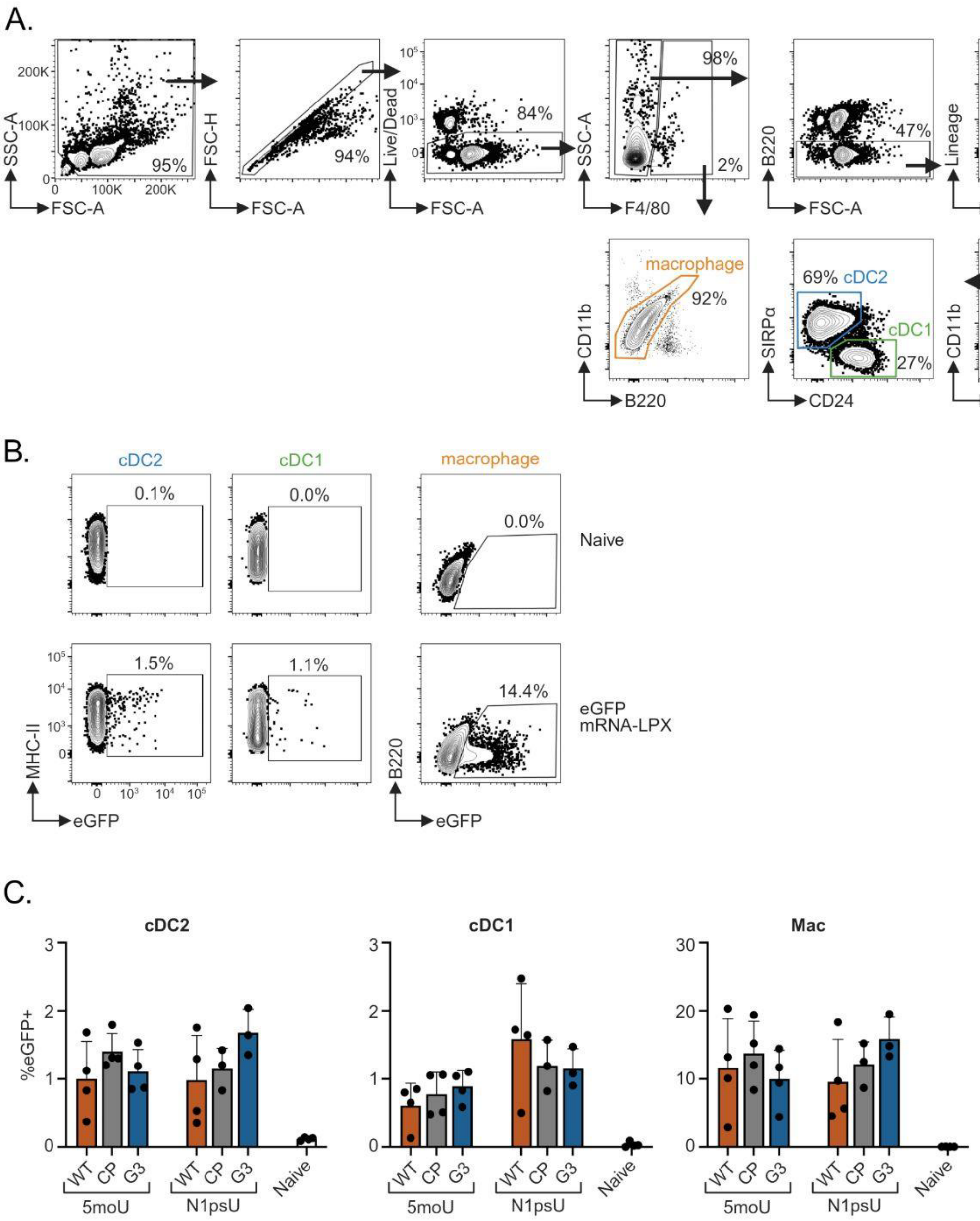


**Supplementary Figure S4. (A)** Representative flow cytometry gating strategy for splenic antigen-presenting cell populations. "Lineage" dump channel includes antibodies against TCRb, CD19, NK1.1, and Ly6G. **(B)** Representative gates for eGFP+ populations. **(C)** Frequency of mRNA-expressing cells (% eGFP+) among splenic populations 24h after injection of 25 µg eGFP mRNA-LPX (i.v.) (n=3-4 per group). Barplots show mean + standard deviation. Statistics from two-way ANOVA with Dunnett's multiple comparisons test, tested against WT T7 for each uridine modification (no significant differences observed with $p<0.05$).

### Supplementary Tables

**Supplementary Table S1:** Quantitative Multi-Attribute Physicochemical Profiling of T7 RNAP Variants Across mRNA Constructs. Values represent the mean percentage of peak area calculated from three independent in vitro transcription (IVT) reactions per construct (n = 3). (a) IP-RPLC Quantification: Tabulated peak area percentages (% pre-peak abortives, % main peak, and % post-peak 3'-loopback hairpins) across eGFP, antigen HiBIT-tagged, and poly(A)-less antigen HiBIT-tagged constructs synthesized using wild-type (WT) T7, commercial polymerase (CP) benchmark, and engineered variants G1–G5. (b) CGE quantification: Tabulated main-peak and abortive peak area percentages measured via denaturing CGE across constructs, orthogonally confirming abortive transcript trends and overall construct integrity. (c) SEC Quantification: Tabulated peak area percentages measured via SEC under non-denaturing conditions, confirming non-covalent aggregate formation for WT T7 and marked aggregate suppression across variants G1–G5 and CP. Highlighted cells (red) indicate significant byproduct accumulation, including elevated abortive pre-peaks (e.g., variant G5), covalent 3'-loopback extensions, or non-covalent aggregates.

a) IP-RPLC Quantification

| mRNA Template | Enzyme | Pre-peak % | Main peak % | Post peak % |
|---|---|---|---|---|
| eGFP | WT T7 | 21.49 | 64.00 | 14.52 |
| | CP | 19.92 | 64.13 | 15.95 |
| | G1 | 23.63 | 60.97 | 15.41 |
| | G2 | 23.55 | 61.69 | 14.76 |
| | G3 | 21.60 | 63.62 | 14.79 |
| | G4 | 19.59 | 65.85 | 14.55 |
| | G5 | 27.58 | 58.68 | 13.74 |
| Antigen HiBIT-tagged construct | WT T7 | 40.50 | 47.33 | 12.18 |
| | CP | 37.57 | 50.26 | 12.18 |
| | G1 | 38.67 | 49.27 | 12.07 |
| | G2 | 39.81 | 48.07 | 12.13 |
| | G3 | 38.10 | 49.25 | 12.66 |
| | G4 | 36.47 | 51.11 | 12.42 |
| | G5 | 40.49 | 47.26 | 12.26 |
| Poly(A)-less antigen HiBIT-tagged construct | WT T7 | 1.16 | 66.39 | 32.45 |
| | CP | 1.44 | 85.89 | 12.68 |

| | | | | |
|---|---|---|---|---|
| | G1 | 2.30 | 84.18 | 13.52 |
| | G2 | 3.69 | 81.79 | 14.53 |
| | G3 | 1.71 | 85.93 | 12.38 |
| | G4 | 1.43 | 86.46 | 12.12 |
| | G5 | 2.00 | 86.23 | 11.77 |

b) CGE quantification

| mRNA Template | Enzyme | Pre-peak % | Main peak % | Post peak % |
|---|---|---|---|---|
| eGFP | WT T7 | 9.20 | 88.20 | 2.65 |
| | CP | 10.05 | 87.55 | 2.35 |
| | G1 | 14.10 | 83.95 | 1.95 |
| | G2 | 21.60 | 76.75 | 1.70 |
| | G3 | 10.65 | 87.15 | 2.15 |
| | G4 | 11.20 | 86.40 | 2.05 |
| | G5 | 20.75 | 77.55 | 1.70 |
| Antigen HiBIT-tagged construct | WT T7 | 28.05 | 68.65 | 3.50 |
| | CP | 33.05 | 64.50 | 2.50 |
| | G1 | 34.75 | 62.50 | 2.70 |
| | G2 | 35.10 | 63.15 | 1.80 |
| | G3 | 32.75 | 65.30 | 2.00 |
| | G4 | 27.70 | 70.45 | 1.90 |
| | G5 | 34.70 | 63.20 | 2.05 |
| Poly(A)-less antigen HiBIT-tagged construct | WT T7 | 8.3 | 69.0 | 22.0 |
| | CP | 5.3 | 92.2 | 2.5 |
| | G1 | 8.6 | 86.7 | 4.6 |
| | G2 | 10.7 | 79.2 | 10.1 |

| | | | | |
|---|---|---|---|---|
| | G3 | 5.3 | 91.9 | 2.8 |
| | G4 | 4.4 | 93.5 | 2.1 |
| | G5 | 8.3 | 88.9 | 2.8 |

c) SEC Quantification

| mRNA Template | Enzyme | Short Impurity % | Intact mRNA % | Aggregates % |
|---|---|---|---|---|
| eGFP | WT T7 | 2.51 | 66.28 | 31.21 |
| | CP | 4.40 | 89.61 | 5.99 |
| | G1 | 5.51 | 87.76 | 6.73 |
| | G2 | 5.60 | 86.92 | 7.47 |
| | G3 | 4.27 | 89.82 | 5.90 |
| | G4 | 4.15 | 90.61 | 5.24 |
| | G5 | 9.24 | 85.38 | 5.38 |
| Antigen HiBIT-tagged construct | WT T7 | 1.71 | 73.3 | 24.99 |
| | CP | 1.74 | 92.56 | 5.71 |
| | G1 | 1.44 | 92.69 | 5.88 |
| | G2 | 1.97 | 92.47 | 5.57 |
| | G3 | 1.25 | 93.59 | 5.18 |
| | G4 | 1.19 | 94.23 | 4.58 |
| | G5 | 1.44 | 93.55 | 5.02 |
| Poly(A)-less antigen HiBIT-tagged construct | WT T7 | 2.82 | 71.24 | 25.95 |
| | CP | 2.04 | 93.04 | 4.93 |
| | G1 | 2.75 | 92.49 | 4.78 |
| | G2 | 3.40 | 87.05 | 9.56 |
| | G3 | 2.15 | 92.75 | 5.11 |

| | | | | |
|---|---|---|---|---|
| | G4 | 2.20 | 92.52 | 5.27 |
| | G5 | 2.57 | 92.23 | 5.21 |

**Supplementary Table S2:** List of anti-mouse antibodies used for the flow cytometry panels.

| Antibody | Clone | Fluorochrome | Supplier | Catalog # | RRID |
|---|---|---|---|---|---|
| B220 | RA3-6B2 | BV605 | BioLegend | 103244 | AB_2563312 |
| CD11b | M1/70 | PerCP-Cy5.5 | BioLegend | 101228 | AB_893232 |
| CD11c | N418 | PE-Cy7 | BioLegend | 117318 | AB_493568 |
| CD19 | 6D5 | AlexaFluor 700 | BioLegend | 115528 | AB_493735 |
| CD24 | M1/69 | BUV395 | BD Biosciences | 744471 | AB_2742259 |
| CD86 | GL-1 | BUV737 | BD Biosciences | 741737 | AB_2871107 |
| F4/80 | BM8 | BV711 | BioLegend | 123147 | AB_2564588 |
| Ly6G | 1A8 | AlexaFluor 700 | BioLegend | 127622 | AB_10643269 |
| MHC-II | M5/114.15.2 | BV421 | BioLegend | 107632 | AB_2650896 |
| SiglecH | 551 | PE | BioLegend | 129606 | AB_2189147 |
| SIRPα | P84 | APC-Fire750 | BioLegend | 144030 | AB_2721317 |
| TCRβ | H57-597 | AlexaFluor 700 | BioLegend | 109224 | AB_1027648 |

**Supplementary Table S3:** List of sgRNA sequences for BMDC KOs.

| **Target Gene** | **Sequence** |
|---|---|
| N/A (NTC) | CGTTAATCGCGTATAATACG |
| N/A (NTC) | CATATTGCGCGTATAGTCGC |
| Ifih1 (MDA5) | CAAGTGGAGACACTCGTCAT |
| Ifih1 (MDA5) | ATGGACGCAGATGTTCGTGG |
| Ddx58 (RIGI) | GTCGGATCTGATAAGACATT |
| Ddx58 (RIGI) | TTGAAAGTCCCAACTTTCGA |
